## Supplemental materials for "The Neural Correlates of Novelty and Variability in Human Decision-Making under an Active Inference Framework"

December 2, 2024

#### Supplementary Material

### 1 Supplementary Method

##### Model-free reinforcement learning

We used a model-free reinforcement learning approach as a baseline model. It involves two value functions, $Q_{Stay/Cue}$ and $Q_{Safe/Risk}$, which respectively describe the expected values of different options for the participants:

$$Q_{Stay/Cue} = \begin{bmatrix} Q_0 & Q_0 \end{bmatrix} \tag{1}$$

$$Q_{Safe/Risk} = \begin{bmatrix} Q_0 & Q_0 \\ Q_0 & Q_0 \end{bmatrix} \tag{2}$$

Here, $Q_0$ equals 6, corresponding to the fixed reward obtained for choosing the safe option. $Q_{Stay/Cue}(1)$ represents the total expected reward for choosing the "Stay" option in this round for the participant, while $Q_{Stay/Cue}(2)$ represents the total expected reward for choosing the "Cue" option in this round. $Q_{Safe/Risk}(1,1)$ and $Q_{Safe/Risk}(1,2)$ respectively denote the expected rewards for choosing the "Safe" option in "Context 1" and "Context 2", whereas $Q_{Safe/Risk}(2,1)$ and $Q_{Safe/Risk}(2,2)$ represent the expected rewards for choosing the "Risk" option in "Context 1" and "Context 2".

$$P_{Stay/Cue}(i) = \frac{e^{\gamma Q_{Stay/Cue}(i)}}{e^{\gamma Q_{Stay/Cue}(1)} + e^{\gamma Q_{Stay/Cue}(2)}} \tag{3}$$

Here, $i = 1$ corresponds to choosing the "Stay" option, and $i = 2$ corresponds to choosing the "Cue" option.

If participants decide whether to choose the "Safe" option or the "Risk" option when knowing the context ("Context 1"):

$$P_{Safe/Risk}(i) = \frac{e^{\gamma Q_{Safe/Risk}(i,1)}}{e^{\gamma Q_{Safe/Risk}(1,1)} + e^{\gamma Q_{Safe/Risk}(2,1)}} \tag{4}$$

If participants decide whether to choose the "Safe" option or the "Risk" option when knowing the context ("Context 2"):

$$P_{Safe/Risk}(i) = \frac{e^{\gamma Q_{Safe/Risk}(i,2)}}{e^{\gamma Q_{Safe/Risk}(1,2)} + e^{\gamma Q_{Safe/Risk}(2,2)}} \tag{5}$$

If participants decide whether to choose the "Safe" option or the "Risk" option when without knowing the context:

$$P_{Safe/Risk}(i) = \frac{e^{\gamma (Q_{Safe/Risk}(i,1) + Q_{Safe/Risk}(i,2))/2}}{\sum_{j=1,2} e^{\gamma (Q_{Safe/Risk}(j,1) + Q_{Safe/Risk}(j,2))/2}} \tag{6}$$

After participants make two choices and receive rewards $r$, $Q_{Stay/Cue}$ and $Q_{Safe/Risk}$ are updated using delta learning rule [1].

If participants chose the "Stay" option:

$$Q_{Stay/Cue}(1) = Q_{Stay/Cue}(1) + \alpha * (r - Q_{Stay/Cue}(1)); \tag{7}$$

If participants chose the "Cue" option:

$$Q_{Stay/Cue}(1) = Q_{Stay/Cue}(1) + \alpha * ((r-1) - Q_{Stay/Cue}(1)); \tag{8}$$

$(-1)$ is the cost of choosing the "Cue" option and $\alpha$ is the learning rate. $Q_{Safe/Risk}(1,1)$ and $Q_{Safe/Risk}(1,2)$ always equal 6 because choosing the "Safe" option consistently results in a fixed reward of 6.

If participants chose the "Risk" option when knowing the context ("Context 1"):

$$Q_{Safe/Risk}(2,1) = Q_{Safe/Risk}(2,1) + \alpha * (r - Q_{Safe/Risk}(2,1)); \tag{9}$$

If participants chose the "Risk" option when knowing the context ("Context 2"):

$$Q_{Safe/Risk}(2,2) = Q_{Safe/Risk}(2,2) + \alpha * (r - Q_{Safe/Risk}(2,2)); \tag{10}$$

If participants chose the "Risk" option when without knowing the context:

$$\begin{aligned} Q_{Safe/Risk}(2,1) &= Q_{Safe/Risk}(2,1) + 0.5 * \alpha * (r - Q_{Safe/Risk}(2,1)); \\ Q_{Safe/Risk}(2,2) &= Q_{Safe/Risk}(2,2) + 0.5 * \alpha * (r - Q_{Safe/Risk}(2,2)). \end{aligned} \tag{11}$$

#### Model-based reinforcement learning

Here we used a model-based reinforcement learning model as another baseline model. It involves the same likelihood matrix mapping from hidden states to outcomes $A$ and the concentration parameters of likelihood $a$.

$$a = \begin{bmatrix} 1 & 1 & prior & prior & 0 & 0 & 0 & 0 \\ 0 & 0 & prior & prior & 0 & 0 & 0 & 0 \\ 0 & 0 & prior & prior & 0 & 0 & 0 & 0 \\ 0 & 0 & prior & prior & 0 & 0 & 0 & 0 \\ 0 & 0 & prior & prior & 0 & 0 & 0 & 0 \\ 0 & 0 & 0 & 0 & 1 & 1 & 0 & 0 \\ 0 & 0 & 0 & 0 & 0 & 0 & 1 & 0 \\ 0 & 0 & 0 & 0 & 0 & 0 & 0 & 1 \end{bmatrix} \tag{12}$$

$$A = Cat(a) \tag{13}$$

When participants decide whether to choose the "Stay" option or the "Cue" option, they consider the expected values of the "Stay" option, the "Cue" option, the "Safe" option, and the "Risk" option under both "Context 1" and "Context 2":

$$Q_{Risk}^{Context1} = A[:,3] \cdot Preference^T, \tag{14}$$

$$Q_{Risk}^{Context2} = A[:,4] \cdot Preference^T, \tag{15}$$

$$Q_{Safe} = Q_{Safe}^{Context1} = Q_{Safe}^{Context2} = 6. \tag{16}$$

Here $Preference$ equals $[6, 12, 9, 3, 0, 0, -1, -1]$. If participants choose the "Stay" option, they will not know the context and make the decision based on the expected values of the "Safe" option and the "Risk" option:

$$Q_{Stay} = max(6, Q_{Risk}^{Context1}/2 + Q_{Risk}^{Context2}/2) \tag{17}$$

If participants choose the "Cue" option, they can then decide whether to choose the "Safe" option or the "Risk" option based on the context:

$$Q_{Cue} = \frac{max(6-1, Q_{Risk}^{Context1} - 1) + max(6-1, Q_{Risk}^{Context2} - 1)}{2} \tag{18}$$

Here $(-1)$ is the cost of choosing the "Cue" option. The probabilities of choosing the "Stay" option and the "Cue" option are respectively:

$$\begin{aligned} P(Stay) &= \frac{e^{\gamma Q_{Stay}}}{e^{\gamma Q_{Stay}} + e^{\gamma Q_{Cue}}}, \\ P(Cue) &= \frac{e^{\gamma Q_{Cue}}}{e^{\gamma Q_{Stay}} + e^{\gamma Q_{Cue}}}. \end{aligned} \tag{19}$$

If participants decide whether to choose the "Safe" option or the "Risk" option when knowing the context ("Context 1"):

$$\begin{aligned} P(Safe) &= \frac{e^{\gamma Q_{Safe}}}{e^{\gamma Q_{Safe}} + e^{\gamma Q_{Risk}^{Context1}}}, \\ P(Risk) &= \frac{e^{\gamma Q_{Risk}^{Context1}}}{e^{\gamma Q_{Safe}} + e^{\gamma Q_{Risk}^{Context1}}}. \end{aligned} \tag{20}$$

If participants decide whether to choose the "Safe" option or the "Risk" option when knowing the context ("Context 2"):

$$\begin{aligned} P(Safe) &= \frac{e^{\gamma Q_{Safe}}}{e^{\gamma Q_{Safe}} + e^{\gamma Q_{Risk}^{Context2}}}, \\ P(Risk) &= \frac{e^{\gamma Q_{Risk}^{Context2}}}{e^{\gamma Q_{Safe}} + e^{\gamma Q_{Risk}^{Context2}}}. \end{aligned} \tag{21}$$

If participants decide whether to choose the "Safe" option or the "Risk" option when without knowing the context:

$$\begin{aligned} P(Safe) &= \frac{e^{\gamma Q_{Safe}}}{e^{\gamma Q_{Safe}} + e^{\gamma (Q_{Risk}^{Context2} + Q_{Risk}^{Context1})/2}}, \\ P(Risk) &= \frac{e^{\gamma (Q_{Risk}^{Context2} + Q_{Risk}^{Context1})/2}}{e^{\gamma Q_{Safe}} + e^{\gamma (Q_{Risk}^{Context2} + Q_{Risk}^{Context1})/2}}. \end{aligned} \tag{22}$$

The process of value updating is the same as the process of value updating in active inference:

$$a = a + \alpha \sum_{\tau} o_\tau \otimes s_\tau. \tag{23}$$

Here, $\alpha$ is the learning rate.

### Supplementary Results

#### Simulation Results

The simulation experiments primarily focus on the utility of AI, AL, and EX in the active inference models, comparing the differences in decision-making among agents with varying AI, AL, and EX parameter values, considering their varying tendencies to avoid risk, reduce ambiguity, and maximize extrinsic rewards. The simulation experiments in the supplementary materials considered two situations $EX = 0$ and $AI = AI = 0$.

When AI and AL are set to 0 and EX is set to 10, it is expected that the agent will solely maximize extrinsic rewards without engaging in exploration. Figure S2 depicts the decisions of such an agent. In the initial trials, the agent almost exclusively selects the “Stay” option, choosing the “Cue” option only occasionally, as the expected rewards for the “Risk” and “Safe” options are both 6 initially, and choosing the “Cue” option involves a cost. Without a bias towards either the “Safe” or “Risk” option, the agent’s second choice is random. In the latter half, as the agent occasionally selects the “Cue” and “Risk” options, updating its expectations of the “Risk” option’s reward, it increases the probability of choosing the “Cue” option. For the second choice, the agent eventually learns the optimal strategy, selecting the “Risk” option in “Context 1” and the “Safe” option in ”Context 2.” While the agent with AI and AL set to 0 can eventually learn the optimal strategy, its rate of strategy optimization is significantly slower compared to an agent with non-zero values for all three parameters.

If EX is set to 0, and AI and AL are set to 10, the agent is expected to solely maximize information gain without pursuing higher extrinsic rewards. Figure S1 illustrates the decisions of such an agent. Regardless of the experiment’s stage, the agent overwhelmingly prefers the “Cue” option in the first choice, as it consistently offers the same value in avoiding risk. Similarly, in the second choice, the agent favors the “Risk” option due to its ability to reduce ambiguity. The agent explores the “Risk” option equally in both “Context 1” and ”Context 2.” However, it is noteworthy that in the latter half of the experiment, the agent’s probability of selecting the “Safe” option increases, as continued exploration of the “Risk” option decreases its perceived ambiguity, rendering its informational value negligible.

In comparison, the active inference model exhibits an excellent exploration mechanism, enabling the agent to quickly learn optimal strategies in uncertain environments. Disabling the model’s AL and AI parameters by setting them to 0 undermines this exploration mechanism, forcing the agent to optimize its strategy slowly through random exploration.

#### Model Recovery

To demonstrate how reliable our models are (the active inference model, model-free reinforcement learning model, and model-based reinforcement learning model), we run some simulation experiments for model recovery. We use these three models, with their own fitting parameters, to generate some simulated data. Then we will fit all three sets of data using these three models. The model recovery results are shown in Fig.S6. This is the confusion matrix of models: the percentage of all subjects simulated based on a certain model that is fitted best by a certain model. The goodness-of-fit was compared using the Bayesian Information Criterion. We can see that the result of model recovery is very good, and the simulated data generated by a model can be best explained by this model.

#### Source Estimate Results

In our results analysis, different criteria lead to different outcomes. For instance, requiring a certain proportion of brain regions to exhibit significant correlations, specifying whether the significance level should be $p < 0.05$ or $p < 0.01$, or mandating a minimum duration for significant correlations in a certain proportion of brain regions all yield distinct results. Therefore, we compiled all brain regions that meet various criteria in the supplementary materials.

#### Supplementary Table

Table S1: In the "Stay/Cue" choice, we require that the activity of more than 50 percent of brain regions remains significantly correlated with the **expected free energy** for over 0.32 seconds, with a significance $p < 0.05$ (after false discovery rate correction). The brain regions are delineated according to the "aparc sub" parcellation.

| **Regressor** | **p value** | **Proportion** | **Duration** | |
|---|---|---|---|---|
| Expected free energy | 0.01 | 0.5 | 0.32s | |
| **Brain Region** | **Duration** | **Proportion** | **regression coefficient** | **t-value** |
| Frontalpole 1-lh | 0.34 | 0.8965 | $1.342 \times 10^{-11}$ | -3.136 |
| Frontalpole 1-rh | 0.372 | 0.9086 | $1.357 \times 10^{-11}$ | -3.228 |
| Lateralorbitofrontal 2-lh | 0.364 | 0.8608 | $1.526 \times 10^{-11}$ | -3.235 |
| Lateralorbitofrontal 3-lh | 0.332 | 0.7691 | $1.362 \times 10^{-11}$ | -3.195 |
| Lateralorbitofrontal 5-lh | 0.336 | 0.7778 | $1.365 \times 10^{-11}$ | -3.126 |
| Lateralorbitofrontal 5-rh | 0.332 | 0.8193 | $1.333 \times 10^{-11}$ | -2.984 |
| Lateralorbitofrontal 6-lh | 0.372 | 0.9086 | $1.550 \times 10^{-11}$ | -3.335 |
| Lateralorbitofrontal 7-lh | 0.332 | 0.8614 | $1.584 \times 10^{-11}$ | -3.062 |
| Lateralorbitofrontal 7-rh | 0.34 | 0.8902 | $1.530 \times 10^{-11}$ | -3.059 |
| Medialorbitofrontal 1-lh | 0.356 | 0.9711 | $1.624 \times 10^{-11}$ | -3.068 |
| Medialorbitofrontal 1-rh | 0.348 | 0.9234 | $1.563 \times 10^{-11}$ | -3.049 |
| Medialorbitofrontal 2-lh | 0.348 | 0.9163 | $1.360 \times 10^{-11}$ | -3.038 |
| Medialorbitofrontal 3-rh | 0.336 | 0.9048 | $1.402 \times 10^{-11}$ | -3.016 |
| Medialorbitofrontal 4-lh | 0.364 | 0.8767 | $1.436 \times 10^{-11}$ | -3.165 |
| Medialorbitofrontal 5-lh | 0.38 | 0.8579 | $1.502 \times 10^{-11}$ | -3.420 |
| Rostralmiddlefrontal 11-lh | 0.364 | 0.8773 | $1.604 \times 10^{-11}$ | -3.200 |
| Rostralmiddlefrontal 11-rh | 0.344 | 0.8469 | $1.486 \times 10^{-11}$ | -3.112 |
| Rostralmiddlefrontal 12-lh | 0.344 | 0.8857 | $1.494 \times 10^{-11}$ | -3.120 |
| Rostralmiddlefrontal 12-rh | 0.336 | 0.7840 | $1.189 \times 10^{-11}$ | -3.210 |
| Rostralmiddlefrontal 13-rh | 0.34 | 0.8510 | $1.428 \times 10^{-11}$ | -3.220 |
| Superiorfrontal 1-lh | 0.336 | 0.9122 | $1.420 \times 10^{-11}$ | -3.101 |

Table S2: In the "Stay/Cue" choice, we require that the activity of more than 50 percent of brain regions remains significantly correlated with the **value of reducing risk** for over 0.152 seconds, with a significance $p < 0.05$ (after false discovery rate correction). The brain regions are delineated according to the "aparc sub" parcellation.

| **Regressor** | **p value** | **Proportion** | **Duration** | |
|---|---|---|---|---|
| Value of avoiding risk | 0.05 | 0.5 | 0.152s | |
| **Brain Region** | **Duration** | **Proportion** | **regression coefficient** | **t-value** |
| Caudalmiddlefrontal 5-lh | 0.152 | 0.9676 | $1.151 \times 10^{-11}$ | -3.363 |
| Caudalmiddlefrontal 6-lh | 0.156 | 0.9837 | $1.006 \times 10^{-11}$ | -3.307 |
| Insula 2-lh | 0.156 | 0.9597 | $1.270 \times 10^{-11}$ | -3.350 |
| Insula 3-lh | 0.156 | 0.9359 | $1.229 \times 10^{-11}$ | -3.251 |
| Medialorbitofrontal 5-lh | 0.164 | 0.8659 | $1.386 \times 10^{-11}$ | -3.081 |
| Parsopercularis 3-lh | 0.160 | 0.9479 | $1.450 \times 10^{-11}$ | -3.334 |
| Postcentral 10-lh | 0.160 | 0.9450 | $1.071 \times 10^{-11}$ | -3.054 |
| Postcentral 11-lh | 0.152 | 0.9737 | $1.111 \times 10^{-11}$ | -3.171 |
| Postcentral 13-lh | 0.152 | 0.8991 | $1.190 \times 10^{-11}$ | -2.951 |
| Precentral 11-lh | 0.152 | 0.9799 | $1.033 \times 10^{-11}$ | -3.376 |
| Precentral 12-lh | 0.156 | 0.9630 | $1.170 \times 10^{-11}$ | -3.280 |
| Precentral 13-lh | 0.152 | 0.9868 | $1.033 \times 10^{-11}$ | -3.347 |
| Precentral 8-lh | 0.152 | 0.9757 | $1.028 \times 10^{-11}$ | -3.434 |
| Precentral 9-lh | 0.152 | 0.9649 | $9.692 \times 10^{-12}$ | -3.323 |
| Rostralanteriorcingulate 2-lh | 0.152 | 0.8816 | $1.007 \times 10^{-11}$ | -2.970 |
| Superiorfrontal 17-lh | 0.152 | 0.9868 | $1.001 \times 10^{-11}$ | -3.371 |
| Supramarginal 3-lh | 0.156 | 0.8521 | $1.260 \times 10^{-11}$ | -3.068 |

Table S3: In the ”Stay/Cue” choice, we require that the activity of more than 50 percent of brain regions remains significantly correlated with the **extrinsic value** for over 0.128 seconds, with a significance $p < 0.05$ (after false discovery rate correction). The brain regions are delineated according to the ”aparc sub” parcellation.

| **Regressor** | **p value** | **Proportion** | **Duration** | |
|---|---|---|---|---|
| Extrinsic value | 0.05 | 0.5 | 0.128s | |
| **Brain Region** | **Duration** | **Proportion** | **regression coefficient** | **t-value** |
| Bankssts 1-rh | 0.136 | 0.9632 | $4.173 \times 10^{-11}$ | 3.547 |
| Bankssts 2-rh | 0.128 | 0.9583 | $3.646 \times 10^{-11}$ | 3.432 |
| Fusiform 7-rh | 0.136 | 0.9647 | $4.727 \times 10^{-11}$ | 3.806 |
| Inferiorparietal 9-rh | 0.132 | 0.9532 | $3.946 \times 10^{-11}$ | 3.560 |
| Inferiortemporal 4-rh | 0.132 | 0.9731 | $5.742 \times 10^{-11}$ | 3.786 |
| Inferiortemporal 5-rh | 0.14 | 0.9771 | $5.648 \times 10^{-11}$ | 3.917 |
| Inferiortemporal 6-rh | 0.136 | 1.0000 | $5.635 \times 10^{-11}$ | 3.651 |
| Inferiortemporal 7-rh | 0.136 | 0.9559 | $4.850 \times 10^{-11}$ | 3.510 |
| Middletemporal 1-rh | 0.132 | 0.9495 | $4.260 \times 10^{-11}$ | 3.326 |
| Middletemporal 3-rh | 0.14 | 0.9679 | $5.060 \times 10^{-11}$ | 3.521 |
| Middletemporal 4-rh | 0.128 | 0.9688 | $4.636 \times 10^{-11}$ | 3.564 |
| Middletemporal 5-rh | 0.132 | 0.9610 | $5.078 \times 10^{-11}$ | 3.490 |
| Middletemporal 6-rh | 0.164 | 0.9338 | $5.983 \times 10^{-11}$ | 3.673 |
| Middletemporal 7-rh | 0.128 | 0.8880 | $4.359 \times 10^{-11}$ | 3.242 |
| Superiortemporal 3-rh | 0.128 | 0.9886 | $3.652 \times 10^{-11}$ | 3.454 |
| Superiortemporal 4-rh | 0.14 | 0.9629 | $4.585 \times 10^{-11}$ | 3.494 |
| Superiortemporal 5-rh | 0.132 | 0.9865 | $4.211 \times 10^{-11}$ | 3.299 |
| Superiortemporal 6-rh | 0.132 | 0.9449 | $3.658 \times 10^{-11}$ | 3.325 |
| Superiortemporal 9-rh | 0.132 | 0.9596 | $3.435 \times 10^{-11}$ | 3.429 |

Table S4: In the result stage after the "Stay/Cue" choice, we require that the activity of more than 50 percent of brain regions remains significantly correlated with **(the value of) avoiding risk** for over 0.32 seconds, with a significance $p < 0.05$ (after false discovery rate correction). The brain regions are delineated according to the "aparc sub" parcellation.

| **Regressor** | **p value** | **Proportion** | **Duration** | |
|---|---|---|---|---|
| (The value of) avoiding risk | 0.05 | 0.5 | 0.32s | |
| **Brain Region** | **Duration** | **Proportion** | **regression coefficient** | **t-value** |
| Caudalanteriorcingulate 1-lh | 0.344 | 0.9341 | $1.523 \times 10^{-11}$ | -3.023 |
| Caudalanteriorcingulate 2-lh | 0.332 | 0.9375 | $1.458 \times 10^{-11}$ | -2.985 |
| Lateralorbitofrontal 1-rh | 0.328 | 0.8659 | $1.733 \times 10^{-11}$ | -2.869 |
| Medialorbitofrontal 5-lh | 0.376 | 0.9109 | $1.839 \times 10^{-11}$ | -3.001 |
| Middletemporal 1-lh | 0.324 | 0.8642 | $2.245 \times 10^{-11}$ | -2.944 |
| Middletemporal 5-lh | 0.344 | 0.9201 | $2.466 \times 10^{-11}$ | -3.098 |
| Parstriangularis 1-rh | 0.364 | 0.9194 | $1.804 \times 10^{-11}$ | -3.103 |
| Parstriangularis 2-rh | 0.344 | 0.8709 | $1.936 \times 10^{-11}$ | -3.038 |
| Parstriangularis 3-rh | 0.328 | 0.9079 | $1.893 \times 10^{-11}$ | -3.117 |
| Parstriangularis 4-rh | 0.364 | 0.9011 | $2.057 \times 10^{-11}$ | -3.278 |
| Rostralanteriorcingulate 2-lh | 0.368 | 0.9348 | $1.553 \times 10^{-11}$ | -3.095 |
| Rostralanteriorcingulate 2-rh | 0.328 | 0.9321 | $1.469 \times 10^{-11}$ | -2.847 |
| Rostralmiddlefrontal 1-rh | 0.344 | 0.9120 | $1.571 \times 10^{-11}$ | -3.043 |
| Rostralmiddlefrontal 10-rh | 0.328 | 0.9011 | $1.772 \times 10^{-11}$ | -3.080 |
| Rostralmiddlefrontal 12-rh | 0.36 | 0.9016 | $1.722 \times 10^{-11}$ | -3.102 |
| Rostralmiddlefrontal 4-rh | 0.348 | 0.8931 | $1.825 \times 10^{-11}$ | -2.994 |
| Rostralmiddlefrontal 5-rh | 0.328 | 0.9463 | $1.664 \times 10^{-11}$ | -3.000 |
| Rostralmiddlefrontal 8-rh | 0.336 | 0.9365 | $1.730 \times 10^{-11}$ | -3.070 |
| Superiorfrontal 5-rh | 0.332 | 0.9277 | $1.548 \times 10^{-11}$ | -2.948 |
| Superiorfrontal 6-rh | 0.328 | 0.9268 | $1.585 \times 10^{-11}$ | -2.966 |
| Superiortemporal 7-lh | 0.34 | 0.8882 | $2.055 \times 10^{-11}$ | -3.017 |

Table S5: In the "Safe/Risk" choice, we require that the activity of more than 90 percent of brain regions remains significantly correlated with the **expected free energy** for over 1.88 seconds, with a significance $p < 0.001$ (after false discovery rate correction). The brain regions are delineated according to the "aparc sub" parcellation.

| **Regressor** | **p value** | **Proportion** | **Duration** | |
|---|---|---|---|---|
| Expected free energy | 0.001 | 0.5 | 1.88s | |
| **Brain Region** | **Duration** | **Proportion** | **regression coefficient** | **t-value** |
| Caudalmiddlefrontal 2-lh | 1.908 | 0.9783 | $2.030 \times 10^{-11}$ | -4.746 |
| Insula 6-lh | 1.896 | 0.9579 | $2.223 \times 10^{-11}$ | -4.693 |
| Middletemporal 5-lh | 1.904 | 0.9850 | $3.350 \times 10^{-11}$ | -5.115 |
| Middletemporal 6-lh | 1.92 | 0.9753 | $3.676 \times 10^{-11}$ | -4.988 |
| Parsorbitalis 2-lh | 1.912 | 0.9187 | $2.694 \times 10^{-11}$ | -4.803 |
| Parstriangularis 1-lh | 1.884 | 0.9581 | $2.580 \times 10^{-11}$ | -4.717 |
| Parstriangularis 2-lh | 1.896 | 0.9768 | $2.619 \times 10^{-11}$ | -4.814 |
| Rostralmiddlefrontal 1-lh | 1.908 | 0.9830 | $2.243 \times 10^{-11}$ | -4.819 |
| Rostralmiddlefrontal 2-lh | 1.896 | 0.9783 | $2.216 \times 10^{-11}$ | -4.789 |
| Rostralmiddlefrontal 4-lh | 1.884 | 0.9662 | $2.082 \times 10^{-11}$ | -4.716 |
| Rostralmiddlefrontal 6-lh | 1.904 | 0.9898 | $2.470 \times 10^{-11}$ | -4.901 |

Table S6: In the "Safe/Risk" choice, we require that the activity of more than 50 percent of brain regions remains significantly correlated with the **value of reducing ambiguity** for over 0.14 seconds, with a significance $p < 0.05$ (after false discovery rate correction). The brain regions are delineated according to the "aparc sub" parcellation.

| **Regressor** | **p value** | **Proportion** | **Duration** | |
|---|---|---|---|---|
| Value of reducing ambiguity | 0.05 | 0.5 | 0.14s | |
| **Brain Region** | **Duration** | **Proportion** | **regression coefficient** | **t-value** |
| Insula 6-lh | 0.144 | 0.9037 | $5.112 \times 10^{-11}$ | -3.165 |
| Lateralorbitofrontal 4-lh | 0.144 | 0.8782 | $4.612 \times 10^{-11}$ | -2.971 |
| Parsorbitalis 2-lh | 0.144 | 0.7381 | $5.591 \times 10^{-11}$ | -3.059 |
| Rostralmiddlefrontal 1-lh | 0.148 | 0.8730 | $4.907 \times 10^{-11}$ | -3.107 |
| Rostralmiddlefrontal 6-lh | 0.16 | 0.8679 | $5.244 \times 10^{-11}$ | -3.067 |
| Superiorfrontal 10-lh | 0.152 | 0.9046 | $4.974 \times 10^{-11}$ | -3.065 |
| Superiorfrontal 6-lh | 0.148 | 0.8845 | $4.779 \times 10^{-11}$ | -2.946 |

Table S7: In the "Safe/Risk" choice, we require that the activity of more than 50 percent of brain regions remains significantly correlated with the **extrinsic value** for over 1.68 seconds, with a significance p $<$ 0.001 (after false discovery rate correction). The brain regions are delineated according to the "aparc sub" parcellation.

| **Regressor** | **p value** | **Proportion** | **Duration** | |
|---|---|---|---|---|
| Extrinsic value | 0.001 | 0.5 | 1.68s | |
| **Brain Region** | **Duration** | **Proportion** | **regression coefficient** | **t-value** |
| Caudalmiddlefrontal 2-lh | 1.712 | 0.9626 | $2.131 \times 10^{-11}$ | 4.607 |
| Caudalmiddlefrontal 3-lh | 1.692 | 0.9485 | $2.081 \times 10^{-11}$ | 4.523 |
| Lateralorbitofrontal 4-lh | 1.688 | 0.9437 | $2.271 \times 10^{-11}$ | 4.595 |
| Middletemporal 4-lh | 1.692 | 0.9063 | $3.402 \times 10^{-11}$ | 4.727 |
| Middletemporal 5-lh | 1.78 | 0.9559 | $3.490 \times 10^{-11}$ | 4.905 |
| Middletemporal 6-lh | 1.764 | 0.9558 | $3.889 \times 10^{-11}$ | 4.847 |
| Middletemporal 6-rh | 1.732 | 0.9301 | $2.865 \times 10^{-11}$ | 4.687 |
| Parsopercularis 2-lh | 1.74 | 0.9478 | $2.697 \times 10^{-11}$ | 4.695 |
| Parsopercularis 4-lh | 1.712 | 0.9528 | $2.647 \times 10^{-11}$ | 4.654 |
| Parsorbitalis 2-lh | 1.748 | 0.9059 | $2.849 \times 10^{-11}$ | 4.675 |
| Parstriangularis 1-lh | 1.732 | 0.9310 | $2.759 \times 10^{-11}$ | 4.655 |
| Parstriangularis 2-lh | 1.76 | 0.9591 | $2.798 \times 10^{-11}$ | 4.739 |
| Parstriangularis 3-lh | 1.704 | 0.9577 | $2.565 \times 10^{-11}$ | 4.673 |
| Precentral 13-lh | 1.696 | 0.9682 | $1.950 \times 10^{-11}$ | 4.744 |
| Precentral 14-lh | 1.74 | 0.9688 | $2.066 \times 10^{-11}$ | 4.799 |
| Rostralmiddlefrontal 1-lh | 1.74 | 0.9664 | $2.351 \times 10^{-11}$ | 4.653 |
| Rostralmiddlefrontal 2-lh | 1.716 | 0.9693 | $2.357 \times 10^{-11}$ | 4.692 |
| Rostralmiddlefrontal 4-lh | 1.712 | 0.9541 | $2.217 \times 10^{-11}$ | 4.634 |
| Rostralmiddlefrontal 5-lh | 1.708 | 0.9653 | $2.333 \times 10^{-11}$ | 4.663 |
| Rostralmiddlefrontal 6-lh | 1.72 | 0.9867 | $2.598 \times 10^{-11}$ | 4.74 |
| Rostralmiddlefrontal 7-lh | 1.7 | 0.9277 | $2.221 \times 10^{-11}$ | 4.602 |
| Rostralmiddlefrontal 8-lh | 1.7 | 0.9788 | $2.520 \times 10^{-11}$ | 4.703 |

Table S8: In the result stage of the "Safe/Risk" choice, we require that the activity of more than 50 percent of brain regions remains significantly correlated with the **extrinsic value** for over 0.248 seconds, with a significance $p < 0.05$ (after false discovery rate correction). The brain regions are delineated according to the "aparc sub" parcellation.

| **Regressor** | **p value** | **Proportion** | **Duration** | |
|---|---|---|---|---|
| Extrinsic value | 0.05 | 0.5 | 0.248s | |
| **Brain Region** | **Duration** | **Proportion** | **regression coefficient** | **t-value** |
| Fusiform 3-rh | 0.252 | 0.9714 | $1.090 \times 10^{-11}$ | -3.202 |
| Fusiform 5-rh | 0.256 | 0.9375 | $9.409 \times 10^{-12}$ | -3.063 |
| Inferiorparietal 11-rh | 0.252 | 0.9538 | $8.761 \times 10^{-12}$ | -3.029 |
| Inferiorparietal 5-rh | 0.256 | 0.9473 | $9.393 \times 10^{-12}$ | -3.067 |
| Inferiortemporal 7-rh | 0.256 | 0.9703 | $1.157 \times 10^{-11}$ | -3.357 |
| Lateraloccipital 3-rh | 0.268 | 0.8806 | $9.523 \times 10^{-12}$ | -2.875 |
| Lateraloccipital 4-rh | 0.26 | 0.9538 | $8.228 \times 10^{-12}$ | -2.902 |
| Lateraloccipital 5-rh | 0.252 | 0.9224 | $9.960 \times 10^{-12}$ | -2.908 |
| Lateraloccipital 8-rh | 0.26 | 0.9790 | $9.969 \times 10^{-12}$ | -3.092 |
| Lateraloccipital 9-rh | 0.252 | 0.9302 | $9.413 \times 10^{-12}$ | -3.026 |
| Lingual 5-rh | 0.256 | 0.9792 | $1.097 \times 10^{-11}$ | -3.155 |
| Paracentral 2-rh | 0.252 | 0.9762 | $5.791 \times 10^{-12}$ | -3.065 |
| Parahippocampal 2-rh | 0.256 | 0.9896 | $1.031 \times 10^{-11}$ | -3.225 |
| Postcentral 10-rh | 0.252 | 0.9707 | $7.185 \times 10^{-12}$ | -2.997 |
| Precuneus 8-rh | 0.256 | 0.9398 | $6.078 \times 10^{-12}$ | -2.904 |
| Superiorparietal 11-rh | 0.26 | 0.9303 | $8.428 \times 10^{-12}$ | -2.895 |
| Superiorparietal 3-rh | 0.268 | 0.9091 | $7.188 \times 10^{-12}$ | -3.0 |
| Superiorparietal 6-rh | 0.264 | 0.9318 | $8.186 \times 10^{-12}$ | -3.01 |
| Superiorparietal 7-rh | 0.256 | 0.9420 | $7.757 \times 10^{-12}$ | -2.958 |
| Superiorparietal 8-rh | 0.264 | 0.9674 | $7.950 \times 10^{-12}$ | -2.989 |

Table S9: In the result stage of the ”Safe/Risk” choice, we require that the activity of more than 50 percent of brain regions remains significantly correlated with **(the value of) reducing ambiguity** for over 0.072 seconds, with a significance $p < 0.05$. The brain regions are delineated according to the ”aparc sub” parcellation.

| **Regressor** | **p value** | **Proportion** | **Duration** | |
|---|---|---|---|---|
| (The value of) reducing ambiguity | 0.05 | 0.5 | 0.072s | |
| **Brain Region** | **Duration** | **Proportion** | **regression coefficient** | **t-value** |
| Paracentral 4-rh | 0.072 | 0.9697 | $2.809 \times 10^{-11}$ | -3.321 |
| Paracentral 5-rh | 0.072 | 0.9506 | $2.803 \times 10^{-11}$ | -3.156 |
| Paracentral 6-rh | 0.072 | 0.9778 | $2.420 \times 10^{-11}$ | -3.145 |
| Precentral 11-rh | 0.072 | 0.9861 | $3.115 \times 10^{-11}$ | -3.316 |
| Precentral 15-rh | 0.072 | 0.9596 | $2.952 \times 10^{-11}$ | -3.278 |
| Precentral 16-rh | 0.072 | 0.9293 | $3.079 \times 10^{-11}$ | -3.23 |
| Precentral 7-rh | 0.072 | 0.9242 | $2.994 \times 10^{-11}$ | -3.27 |
| Superiorparietal 3-rh | 0.072 | 0.9343 | $2.940 \times 10^{-11}$ | -3.132 |
| Superiorparietal 6-rh | 0.072 | 0.9611 | $3.410 \times 10^{-11}$ | -3.19 |
| Supramarginal 1-rh | 0.072 | 0.9615 | $3.942 \times 10^{-11}$ | -3.41 |
| Supramarginal 9-rh | 0.072 | 0.9213 | $3.731 \times 10^{-11}$ | -3.452 |

### Supplementary Figure

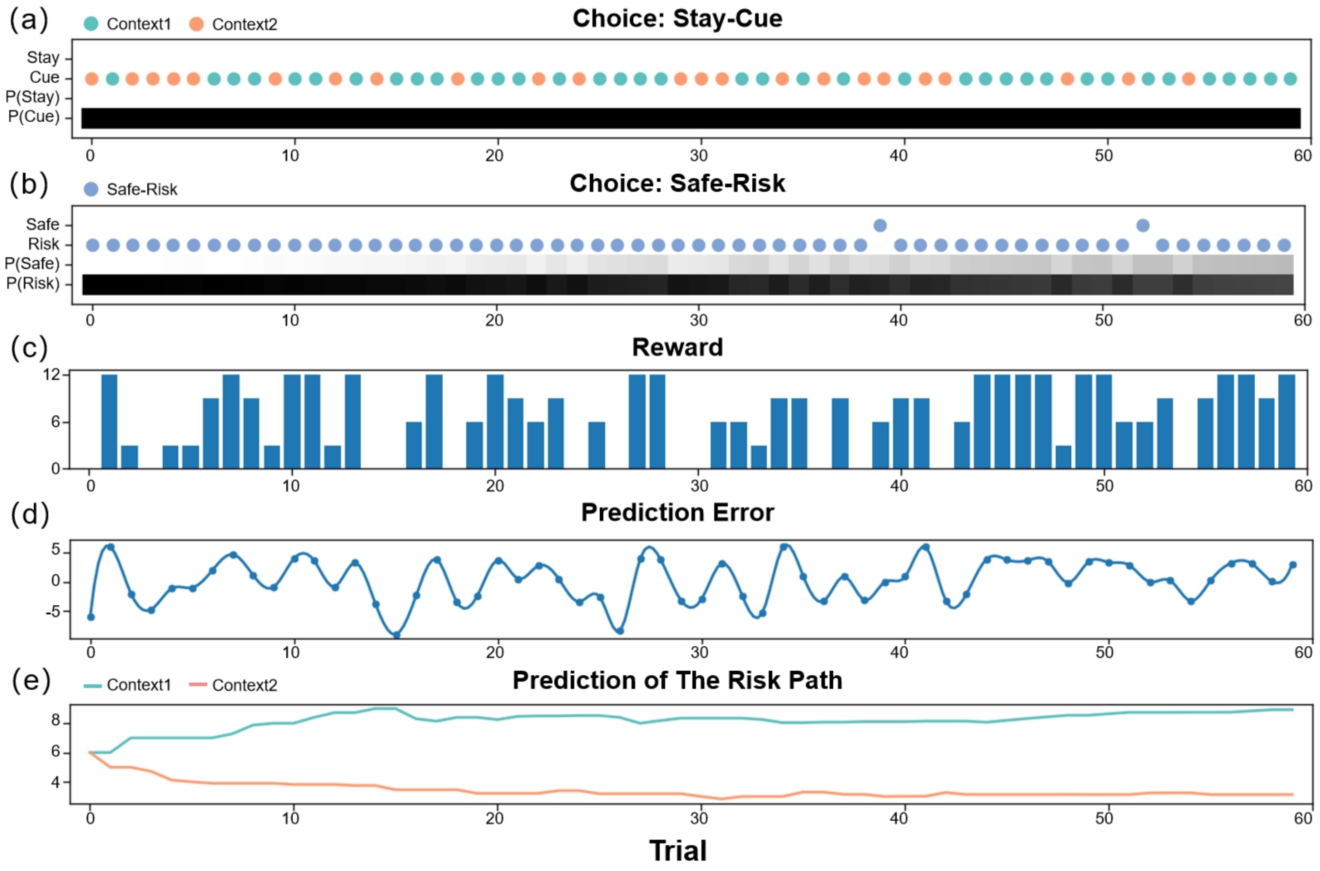


Figure S1: The simulation experiment results. This figure demonstrates how an agent selects actions and updates beliefs over 60 trials in the active inference framework. The first two panels (a-b) display the agent's policy and depict how the policy probabilities are updated (choosing between the stay or cue option in the first choice, and selecting between the safe or risky option in the second choice). The scatter plot indicates the agent's actions, with green representing the cue option when the context of the risky path is "Context 1" (high-reward context), orange representing the cue option when the context of the risky path is "Context 2" (low-reward context), purple representing the stay option when the agent is uncertain about the context of the risky path, and blue indicating the safe-risky choice. The shaded region represents the agent's confidence, with darker shaded regions indicating greater confidence. The third panel(c) displays the rewards obtained by the agent in each trial. The fourth panel(d) shows the prediction error of the agent in each trial. Finally, the fifth panel(e) illustrates the expected rewards of the "Risky Path" in the two contexts of the agent.

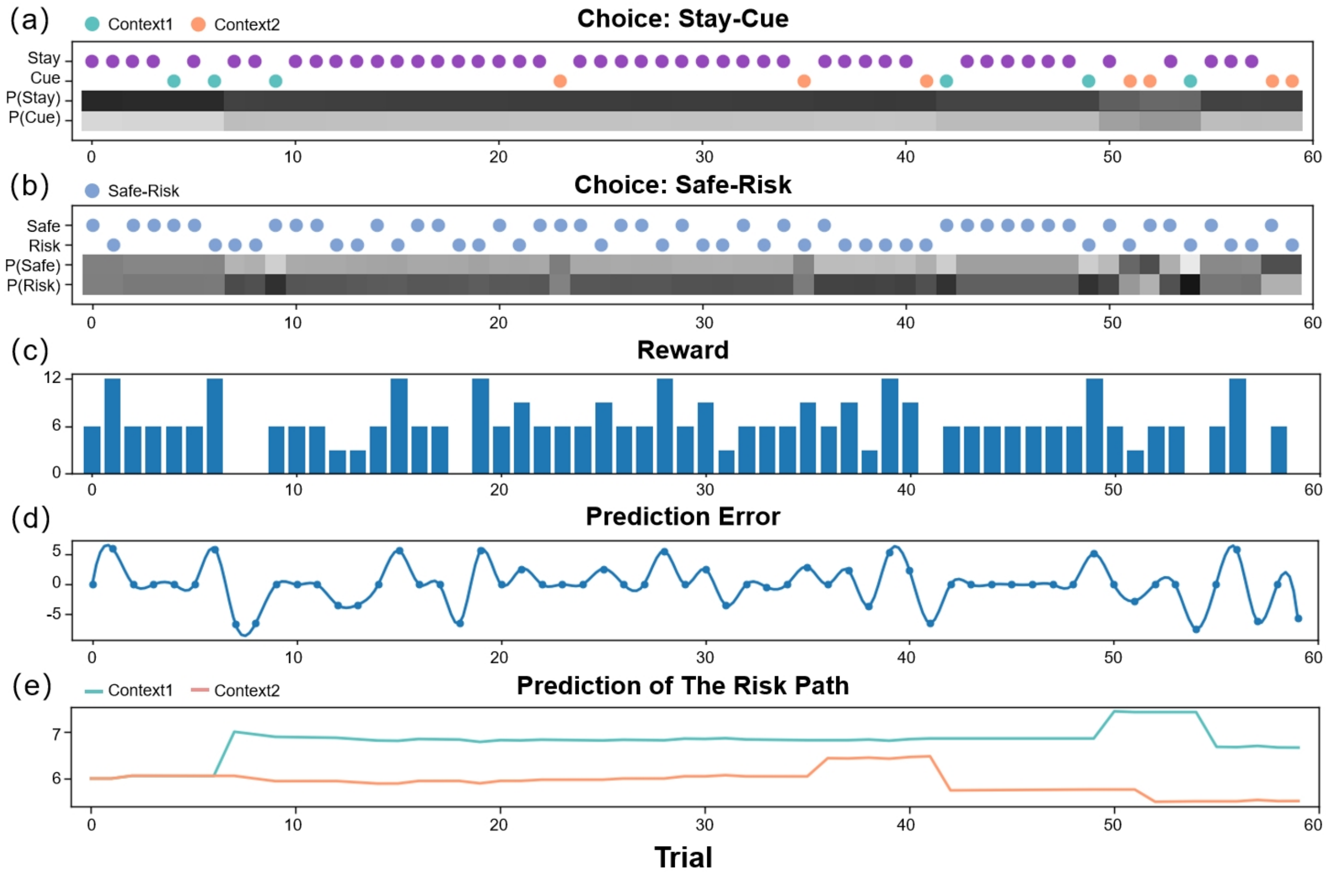


Figure S2: The simulation experiment results. This figure demonstrates how an agent selects actions and updates beliefs over 60 trials in the active inference framework. The first two panels (a-b) display the agent's policy and depict how the policy probabilities are updated (choosing between the stay or cue option in the first choice, and selecting between the safe or risky option in the second choice). The scatter plot indicates the agent's actions, with green representing the cue option when the context of the risky path is "Context 1" (high-reward context), orange representing the cue option when the context of the risky path is "Context 2" (low-reward context), purple representing the stay option when the agent is uncertain about the context of the risky path, and blue indicating the safe-risky choice. The shaded region represents the agent's confidence, with darker shaded regions indicating greater confidence. The third panel(c) displays the rewards obtained by the agent in each trial. The fourth panel(d) shows the prediction error of the agent in each trial. Finally, the fifth panel(e) illustrates the expected rewards of the "Risky Path" in the two contexts of the agent.

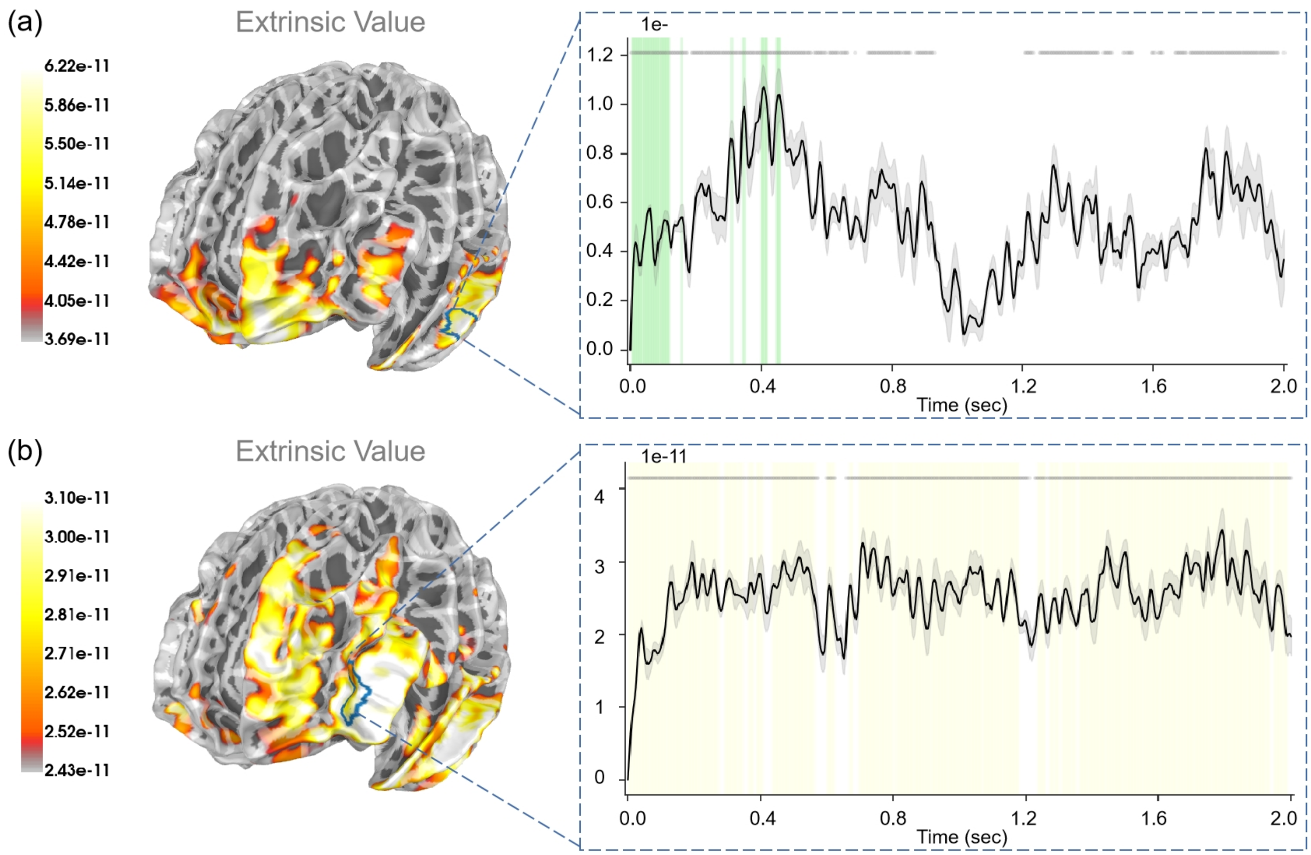


Figure S3: The source estimation results of extrinsic value in the two choosing stages. (A) The regression intensity ($\beta$) of extrinsic value in the "First choice" stage. The right panel indicates the regression intensity between the middle temporal gyrus (6, right half) and extrinsic value. The green-shaded regions indicate $p < 0.05$ after FDR correction (the average $t-value$ during these significant periods equals 3.673). (B) The regression intensity ($\beta$) of extrinsic value in the "Second choice" stage. The right panel indicates the regression intensity between the rostral middle frontal gyrus (6, left half) and extrinsic value. The yellow-shaded regions indicate $p < 0.001$ after FDR correction (the average $t-value$ during these significant periods equals 4.740). The black lines indicate the average intensities and the gray-shaded regions indicate the ranges of variations. The gray lines indicate $p < 0.05$ before FDR.

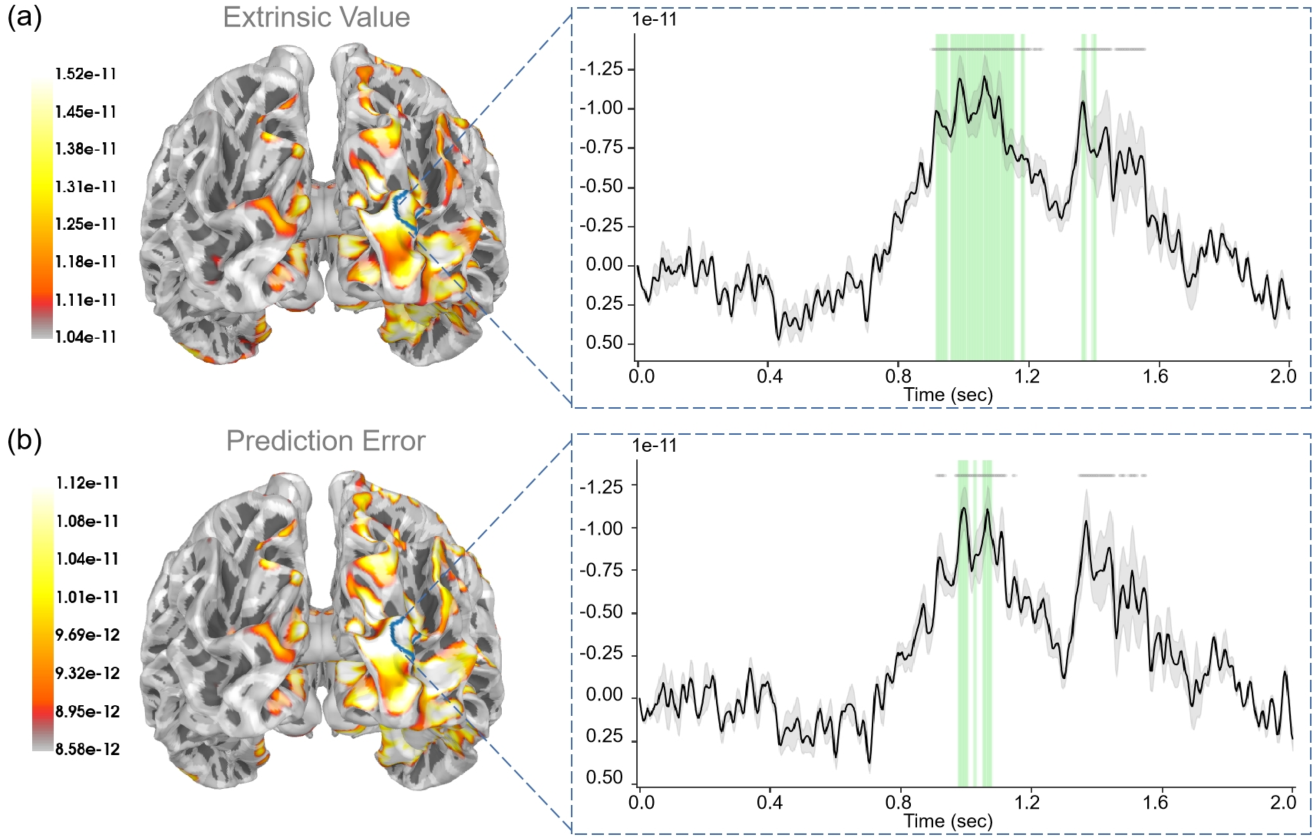


Figure S4: The source estimation results of extrinsic value and prediction error in the "Second result" stage. (A) The regression intensity ($\beta$) of extrinsic value. The right panel indicates the regression intensity between the lateral occipital cortex (3, right half) and extrinsic value. The green-shaded regions indicate $p < 0.05$ after FDR correction (the average $t - value$ during these significant periods equals 2.875). (B) The regression intensity ($\beta$) of prediction error. The right panel indicates the regression intensity between the lateral occipital cortex (3, right half) and prediction error. The green-shaded regions indicate $p < 0.05$ after FDR correction (the average $t - value$ during these significant periods equals $-2.716$). The black lines indicate the average intensities and the gray-shaded regions indicate the ranges of variations. The gray lines indicate $p < 0.05$ before FDR.

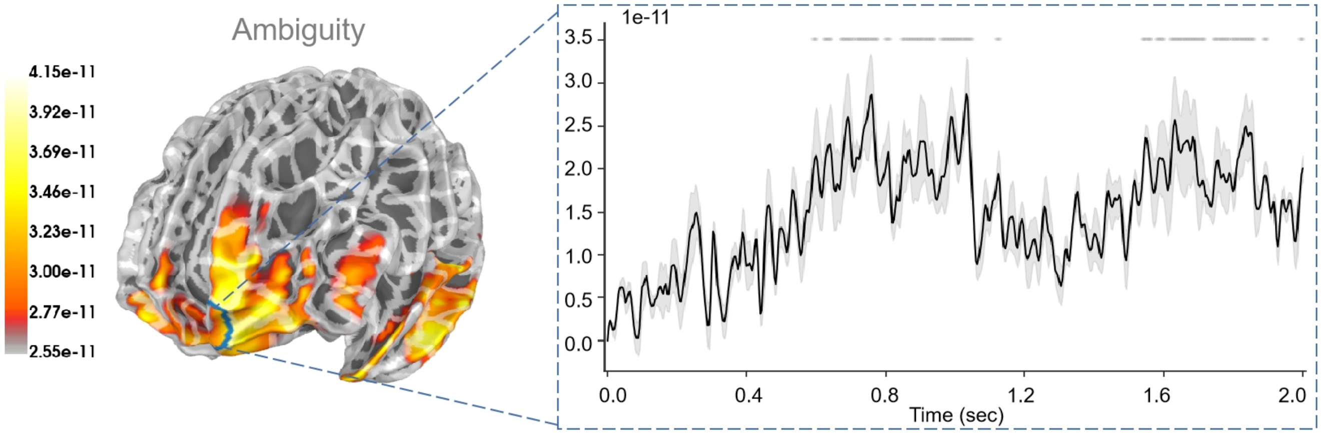


Figure S5: The source estimation results of ambiguity in the "Second choice" stage. The right panel indicates the regression intensity between the frontal pole (1, left half) and ambiguity. The black line indicates the average intensities and the gray-shaded regions indicate the ranges of variations. The gray lines indicate $p < 0.05$ before FDR.

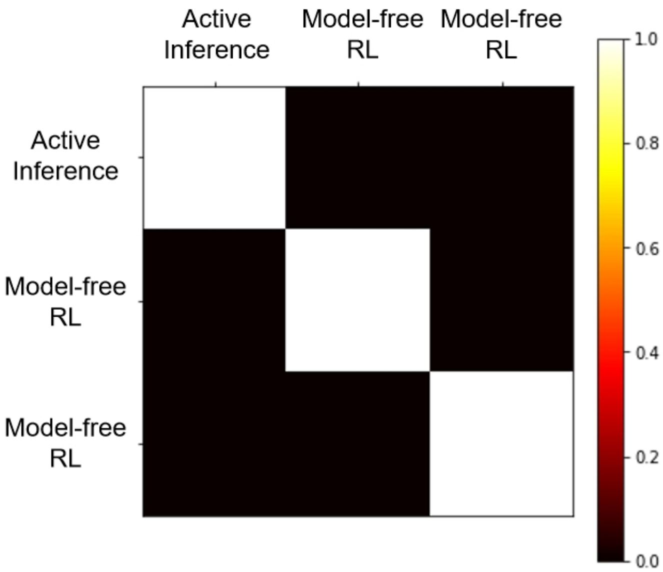


Figure S6: Model Recovery Results.
